# A novel human fetal liver-derived model reveals that MLL-AF4 drives a distinct fetal gene expression program in infant ALL

**DOI:** 10.1101/2020.11.15.379990

**Authors:** Siobhan Rice, Thomas Jackson, Nicholas T Crump, Nicholas Fordham, Natalina Elliott, Sorcha O’Byrne, Sarah Inglott, Dariusz Ladon, Gary Wright, Jack Bartram, Philip Ancliff, Adam J Mead, Christina Halsey, Irene Roberts, Thomas A Milne, Anindita Roy

## Abstract

Although 90% of children with acute lymphoblastic leukemia (ALL) are now cured^1^, the prognosis of infant-ALL (diagnosis within the first year of life) remains dismal^2^. Infant-ALL is usually caused by a single genetic hit that arises *in utero*: rearrangement of the *MLL/KMT2A* gene (*MLL-r*). This is sufficient to give rise to a uniquely aggressive and treatment-refractory leukemia compared to older children with the same *MLL-r*^3–5^. The reasons for disparate outcomes in patients of different ages with identical driver mutations are unknown. This paper addresses the hypothesis that fetal-specific gene expression programs co-operate with MLL-AF4 to initiate and maintain infant-ALL. Using direct comparison of fetal and adult HSC and progenitor transcriptomes we identify fetal-specific gene expression programs in primary human cells. We show that *MLL-AF4*-driven infant-ALL, but not *MLL-AF4* childhood-ALL, displays expression of fetal-specific genes. In a direct test of this observation, we find that CRISPR-Cas9 gene editing of primary human fetal liver cells to produce a t(4;11)/*MLL-AF4* translocation replicates the clinical features of infant-ALL and drives infant-ALL-specific and fetal-specific gene expression programs. These data strongly support the hypothesis that fetal-specific gene expression programs co-operate with MLL-AF4 to initiate and maintain the distinct biology of infant-ALL.

## MAIN

In >70% of infant-ALL cases, the main driver mutation is a chromosomal translocation that leads to rearrangement of the *Mixed Lineage Leukemia* (*MLL/KMT2A*) gene (*MLL-r*)^2,6,7^ producing MLL fusion proteins such as MLL-AF4^8^. MLL-AF4 binds directly to gene targets where it aberrantly upregulates gene expression, partly by increasing histone-3-lysine-79 dimethylation (H3K79me2) through DOT1L recruitment^9^. The prevalence of *MLL-r* in infant ALL contrasts with what is observed in childhood-ALL, where *MLL-r* accounts for only 2-5%of cases^3,10^. Intriguingly, *MLL-r* childhood-ALL has an event-free survival (EFS) of 50-59%^3,4,11^ compared to 19-45% in *MLL-r* infant-ALL^4,5^. This inferior outcome for *MLL-r* infant-ALL does not appear to be due to age-related differences in drug metabolism and/or toxicity since *MLL* wild-type (*MLL*wt) infant-ALL has excellent EFS (74-93%)^7,12^. This suggests there may be intrinsic biological differences between *MLL-r* infant-ALL and *MLL-r* childhood-ALL blasts. In support of this, the *MLL* breakpoint region tends to differ in *MLL-r* infant-ALL^13^ compared to *MLL-r* childhood-ALL, and infant-ALL is associated with a high frequency of the poor prognosis *HOXA^lo^/IRX^hi^ MLL-r* molecular profile^14^. However, very little is known about the underlying reasons for these age-related differences.

A characteristic and baffling feature of *MLL-r* infant-ALL is the fact that this single hit before birth seems to be sufficient to induce a rapidly-proliferating therapy-resistant leukemia without the need for additional mutations^15^, unlike many cases of childhood-ALL which also originate *in utero* but only develop into full-blown leukemia after a second post-natal hit^15^. One reason for this could be that the specific fetal progenitors in which the translocation arises provide the permissive cellular context necessary to cooperate with *MLL-r* to induce infant-ALL^16–18^.

To investigate this, we used the most common *MLL-r* infant-ALL, *MLL-AF4*, as a disease model^8^. Using a previously published patient bulk RNA-seq dataset containing both *MLL-AF4* infant-ALL (n=19) and *MLL-AF4* childhood-ALL (n=5) samples^15^, we found that *MLL-AF4* childhood-ALL clustered separately from *MLL-AF4* infant-ALL (Fig. 1a). In addition, we observed 2 sub-clusters of *MLL-AF4* infant-ALL, representing *HOXA^hi^/IRX^lo^* and *HOXA^lo^/IRX^hi^* infant-ALL subsets, which have previously been characterized (Fig. 1a, Supplementary Figs. 1a and 1b)^14^. Differential gene expression analysis between *MLL-AF4* infant-ALL and *MLL-AF4* childhood-ALL identified 617 significantly differentially expressed genes (FDR < 0.05), 193 of which were upregulated in *MLL-AF4* infant-ALL and therefore represented an infant-ALL-specific gene expression profile (Fig. 1b, Supplementary Fig. 1c, Supplementary Table 1). The two most significantly upregulated genes in *MLL-AF4* infant-ALL were *HOXB4* and *HOXB3* (Fig. 1c, Supplementary Fig. 1d). These *HOXB* genes were effective as a marker of infant-ALL regardless of the *HOXA/IRX* status of the infant-ALL patients (Fig. 1d).

**Fig. 1.**
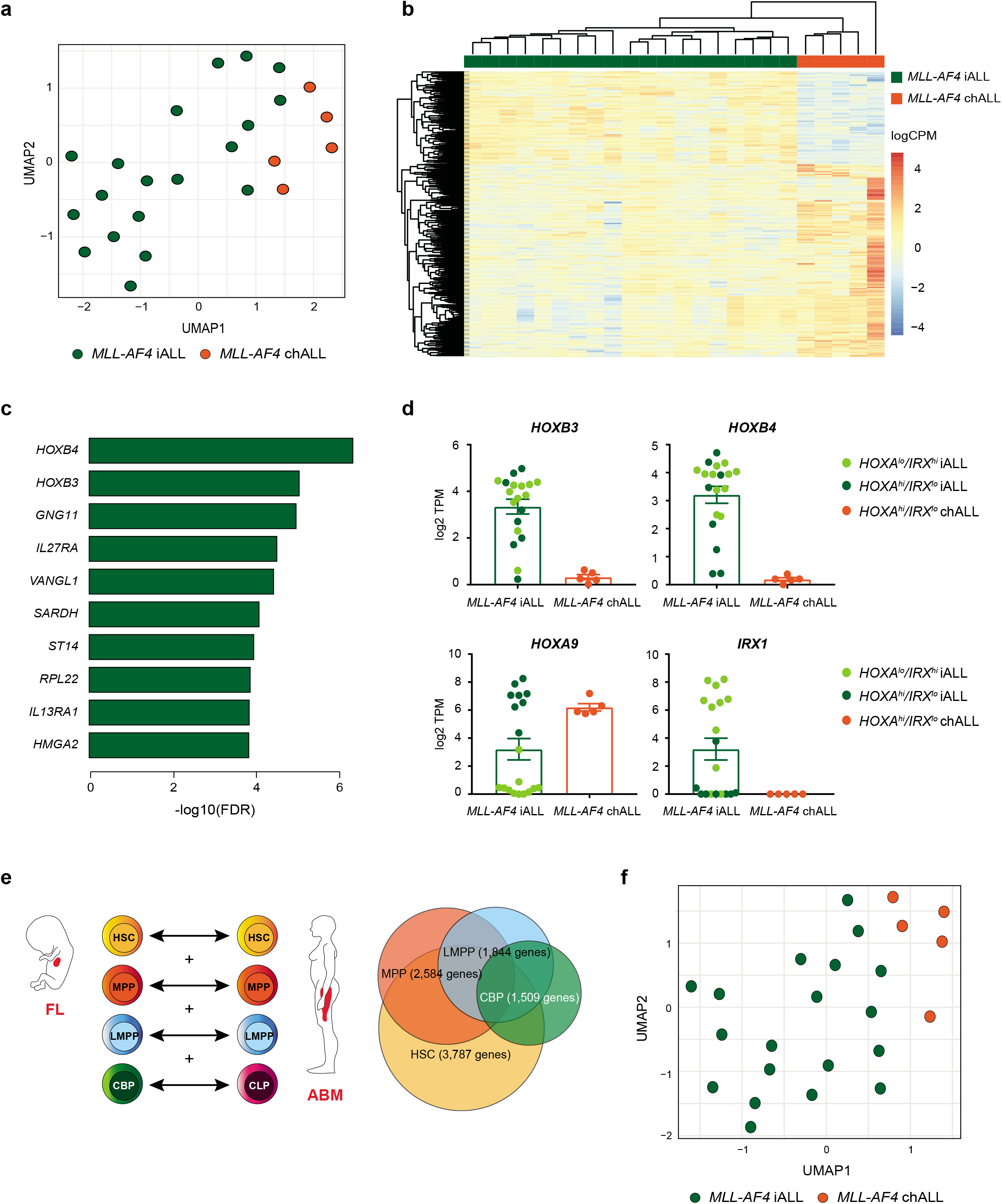
Fetal gene expression programs drive the distinct molecular profile of *MLL-AF4* infant-ALL (iALL) **a.** UMAP showing clustering of *MLL-AF4* infant-ALL (iALL (green), n=19) and *MLL-AF4* childhood-ALL (chALL (orange), n=5) from a previously published patient dataset^15^ based on the 500 most variable genes. **b.** Heatmap showing clustering of *MLL-AF4* infant-ALL (iALL (green), n=19) and *MLL-AF4* childhood-ALL (chALL (orange), n=5) based on 617 significantly differentially expressed genes (FDR<0.05, Supplementary Table 1). Color scale = log2 counts per million (logCPM) **c.** Barplot showing significance (−log10(FDR)) for the 10 most significantly upregulated genes in *MLL-AF4* infant-ALL. **d.** Expression of *HOXB3, HOXB4, HOXA9* and *IRX1* in *MLL-AF4* infant-ALL (light green = *HOXA^lo^/IRX^hi^* infant-ALL (iALL), n=11; dark green = *HOXA^hi^/IRX^lo^* infant-ALL (iALL), n=8 (see Supplementary Fig. 1a and 1b)) and *MLL-AF4* childhood-ALL (*HOXA^hi^/IRX^lo^* chALL, orange, n=5). Individual values are given as log2 transcripts per million (TPM). Data shown as mean ± SEM. **e.** (left) Schematic representation of differential gene expression analysis between FL and ABM. Equivalent HSPC subpopulations were compared and significantly differentially expressed genes (FDR<0.05) for all 4 comparisons were combined into a master list of genes that were differentially expressed in at least 1 HSPC subpopulation (HSC = hematopoietic stem cell, MPP = multipotent progenitor cell, LMPP = lymphoid-primed multipotent progenitor cell, CBP = committed B progenitor, CLP = common lymphoid progenitor). (right) Venn diagram showing overlap of differentially expressed genes for each HSPC subpopulation (see Supplementary Table 2). **f.** UMAP showing clustering of *MLL-AF4* infant-ALL (iALL (green), n=19) and *MLL-AF4* childhood-ALL (chALL (orange), n=5) from a previously published patient dataset^15^ based on 5,709 genes differentially expressed between FL HSPCs and ABM HSPCs (see Supplementary Table 2).

We next sought to determine the extent to which normal fetal gene expression programs contribute to the distinct molecular profile of *MLL-AF4* infant-ALL. We compared bulk RNA-seq for sorted human fetal liver (FL) hematopoietic stem and progenitor cell (HSPC) subpopulations previously generated in our lab^19^ to a human adult bone marrow (ABM) HSPC RNA-seq dataset^20^. We carried out differential gene expression analysis between comparable subpopulations of FL and ABM HSPCs along the B lineage differentiation pathway (Fig. 1e). The hematopoietic stem cell (HSC) subpopulation shows the greatest number of differentially expressed genes between FL and ABM (3,787 genes), reducing at each subsequent stage of B lineage differentiation (1,509 genes differentially expressed between FL committed B progenitors (CBP) and ABM common lymphoid progenitors (CLP)) (Fig. 1e). A total of 5,709 genes were differentially expressed between FL and ABM in a least one HSPC subpopulation when we combined all differentially expressed gene lists (Fig. 1e, Supplementary Table 2).

We repeated the clustering analysis of the patient dataset based on these 5,709 genes and found that they were capable of separating *MLL-AF4* infant-ALL from *MLL-AF4* childhood-ALL (Fig. 1f, Supplementary Fig. 2a). Comparing differentially expressed genes in both the normal and leukemic setting, we found 72 genes that were significantly upregulated in both normal FL HSPCs and *MLL-AF4* infant-ALL (~40% of all genes upregulated in *MLL-AF4* infant-ALL compared to *MLL-AF4* childhood-ALL) (Supplementary Fig. 2b, Supplementary Table 2), including *IGF2BP1*, a member of the fetal-specific *LIN28B* gene expression pathway^21^, which has previously been reported to positively regulate *HOXB4* expression^22^ (Supplementary Fig. 2c). Together, these data suggest that the molecular profile of the human fetal HSPCs that form the target cells for leukemic transformation plays a role in determining the distinct gene expression profile of *MLL-AF4* infant-ALL.

To test the hypothesis that to accurately model *MLL-AF4* infant-ALL, the *MLL-AF4* translocation should be expressed in human fetal HSPCs, we directly induced the most common t(4;11)/*MLL-AF4* translocation in infant-ALL (with the *MLL* breakpoint in intron 11^13^) in 13-15 post-conception week (pcw) human FL CD34+ cells by CRISPR-Cas9 genome editing. Edited samples (n=3) and biologically-matched mock-edited controls (n=3) were transferred to MS-5 co-cultures to facilitate expansion of successfully edited cells along the B lineage (designated *^CRISPR^MLL-AF4+*).

By week 3 of co-culture, CD19+ B cell numbers were >900-fold higher in *^CRISPR^MLL-AF4+* cultures compared to controls (p<0.005), suggesting that the translocation had successfully transformed the cells (Fig. 2a). RT-qPCR confirmed expression of both *MLL-AF4* and *AF4-MLL* in *^CRISPR^MLL-AF4+* cells but not controls (Fig. 2b). Virtually all human cells generated from *^CRISPR^MLL-AF4+* cultures (Supplementary Fig. 3b) were CD19+ B cells, compared to <20% in control cultures (Fig. 2c and 2d). Although there were fewer residual CD34+ cells in *^CRISPR^MLL-AF4+* cultures, the majority of these were CD19+ B progenitors, unlike control CD34+ cells, suggesting that *MLL-AF4*-driven B lineage specification occurs at a progenitor stage (Fig. 2d, right). More detailed immunophenotyping showed that the majority of *^CRISPR^MLL-AF4+* cells were CD34^-^CD19^+^CD10^+^IgM/IgD^-^ preB cells, of which ~10% aberrantly expressed the leukemia-associated marker CD133, a direct gene target of MLL-AF4^23^ (Figs. 2c and 2d). By week 7 of co-culture, when control cultures no longer produced any detectable human cells, the number of human cells in *^CRISPR^MLL-AF4+* cultures began to decline (Fig. 2a), suggesting MS-5 stroma may not be optimal for long-term maintenance of FL-derived *^CRISPR^MLL-AF4+* cells.

**Fig. 2.**
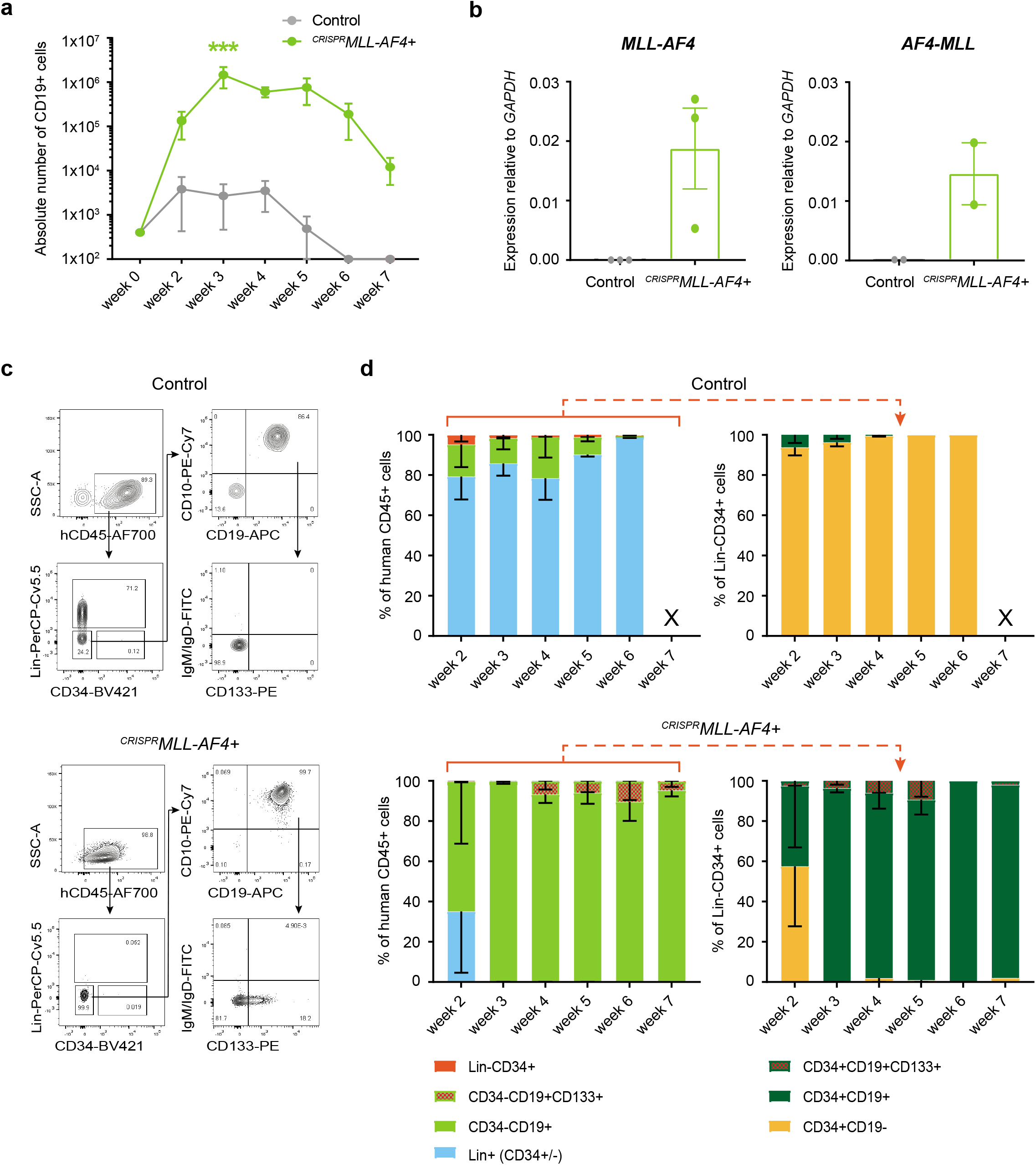
A CRISPR-Cas9-induced t(4;11) *MLL-AF4* translocation in human FL HSPCs causes a dramatic increase in B cell proliferation *in vitro*. **a.** Cumulative absolute number of human CD45+CD19+ cells per well over time during *^CRISPR^MLL-AF4+* and control MS-5 co-culture (n=3). *** =p<0.005 (Two-way ANOVA with Sidak correction for multiple comparisons). Data shown as mean ± SEM. **b.** RT-qPCR of human CD45+ cells showing expression of *MLL-AF4* (n=3) and *AF4-MLL* (n=2) relative to *GAPDH* at week 4 of MS-5 co-culture. Data shown as mean ± SEM. **c.** Representative flow cytometry plots of viable, single cells from control and *^CRISPR^MLL-AF4+* cultures on week 4 of co-culture. Custom lineage cocktail (Lin) = CD2/CD3/CD14/CD16/CD56/CD235a (see Supplementary Table 5). **d.** Quantification of human cell immunophenotypes as a percentage of human CD45+ (hCD45+) cells (left), and progenitor immunophenotypes as a percentage of hCD45+Lin-CD34+ cells (right), in control (n=2-3) and *^CRISPR^MLL-AF4+* (n=2-3) cultures over time. X = data not shown as total number of hCD45+ cells < 50. Data shown as mean ± SEM.

To test whether *^CRISPR^MLL-AF4+* cells could generate leukemia *in vivo*, human FL CD34+ cells (13 pcw, n=4) were edited as before and transplanted into sub-lethally irradiated NSG mice (*^CRISPR^MLL-AF4+*, n=3; control, n=5). By 12 weeks post-transplant, human CD45+ cells were detected in peripheral blood (PB) (Supplementary Fig. 4a) and RT-qPCR showed that both *MLL-AF4* and *AF4-MLL* fusion transcripts were clearly detectable in human CD45+ cells from *^CRISPR^MLL-AF4+* mice (Supplementary Fig. 4b). B-ALL rapidly developed in all three *^CRISPR^MLL-AF4+* mice with median latency of 18 weeks, whereas no control mice (0/5) developed any form of leukemia (Fig. 3a). FISH analysis (Fig 3b) and Sanger sequencing (Supplementary Fig. 4c) confirmed the presence of a heterozygous *MLL-AF4* translocation in *^CRISPR^MLL-AF4+* cells.

**Fig 3.**
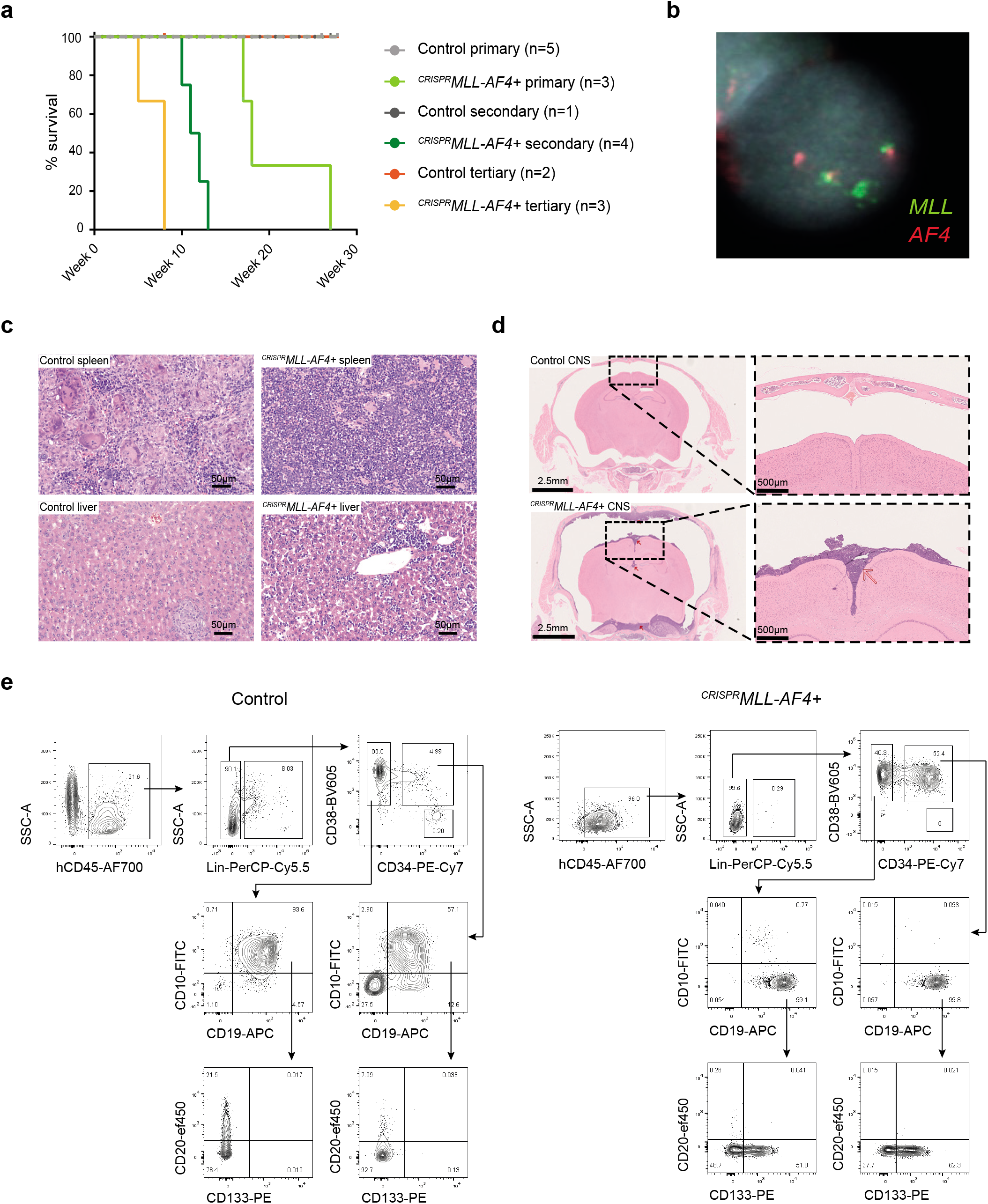
*^CRISPR^MLL-AF4+* cells give rise to a B-ALL *in vivo* that recapitulates many key features of iALL. **a.** Leukemia-free survival for primary (*^CRISPR^MLL-AF4+* n=3; control n=5), secondary (*^CRISPR^MLL-AF4+* n=4; control n=1) and tertiary (*^CRISPR^MLL-AF4+* n=3; control n=2) recipient mice. Mice culled with no signs of leukemia (see Online Methods) are censored (shown as tick above line). Latency significantly reduced for secondary (p<0.02) and tertiary (p<0.03) *^CRISPR^MLL-AF4+* compared to primary *^CRISPR^MLL-AF4+* (Log-Rank (Mantel-Cox) test). **b.** Dual FISH (*MLL* probe = green; *AF4* probe = red) showing heterozygous chromosomal translocation in a *^CRISPR^MLL-AF4+* cell isolated from the spleen of a primary recipient mouse. Representative image of 200 cells analyzed. **c.** Representative H&E staining of spleen and liver from control and *^CRISPR^MLL-AF4+* primary recipient mice. Scale bar = 50μm. **d.** Representative H&E staining of control and *^CRISPR^MLL-AF4+* primary recipient mouse heads (scale bar = 2.5mm; red arrows = regions of concentrated blast cell infiltration). High magnification images (scale bar = 500μm) highlight striking parameningeal blast cell infiltration in *^CRISPR^MLL-AF4+* mice (red arrow) but not control. **e.** Representative flow cytometry plots of viable, single cells in control and *^CRISPR^MLL-AF4+* BM at termination (week 17 and 18 respectively).

The B-ALL in *^CRISPR^MLL-AF4+* mice recapitulated key phenotypic features of infant-ALL, including circulating blasts in the PB (Supplementary Fig. 4d), and blast infiltration into spleen and liver (Fig. 3c, Supplementary Fig. 4e)^24–26^. *^CRISPR^MLL-AF4+* mice also had central nervous system (CNS) disease, with extensive parameningeal blast cell infiltration (Fig. 3d); a key clinical feature of infant-ALL that has not been previously reported in *MLL-AF4* mouse models^27–29^. Although the clinico-pathological features were the same in all leukemic mice (Supplementary Table 3), 2/3 *^CRISPR^MLL-AF4+* mice had a CD19+CD10-CD20-IgM/IgD-CD34+/- proB ALL immunophenotype (Fig. 3e, Supplementary Fig. 4f), while the remaining mouse had a preB ALL immunophenotype, with majority of the cells being CD19+CD10+CD20-IgM/IgD-CD34+/- (Supplementary Fig. 4f and 4g). Further characterization of *^CRISPR^MLL-AF4+* proB ALL revealed that it recapitulated the immunophenotype of *MLL-AF4* infant-ALL, including heterogeneous expression of CD133^23^, NG2^30^ and CD24 (Fig. 3e, Supplementary Fig. 4h). Sequencing of the IgH locus showed that *^CRISPR^MLL-AF4+* ALL was clonal (Supplementary Table 3).

Secondary (n=4) and tertiary (n=3) recipient mice all developed B-ALL with significantly reduced latency compared to primary recipients (median survival 11.5 weeks in secondary (p<0.02) and 8 weeks in tertiary (p<0.03)) (Fig. 3a). The clinico-pathological and immunophenotypic features of primary *^CRISPR^MLL-AF4+* ALL were maintained in secondary recipients, including CNS disease (Supplementary Table 3). Together these data show that CRISPR-Cas9 induced *MLL-AF4* translocation in human FL is sufficient to promote a rapidly progressive, fatal, transplantable B-ALL that recapitulates key features of infant-ALL.

We compared bulk RNA-seq from control and *^CRISPR^MLL-AF4+* bone marrow (Supplementary Fig. 5a) to two independent patient datasets^15,19^ and found that, on a transcriptome-wide level, *^CRISPR^MLL-AF4+* ALL more closely resembled *MLL-AF4* ALL patients compared to *MLL*wt ALL patients (Fig. 4a^19^ and 4b, Supplementary Fig. 5b^15^). Moreover, *^CRISPR^MLL-AF4+* ALL resembled *HOXA^lo^/IRX^hi^ MLL-AF4* infant-ALL (Supplementary Fig. 5c)^14^. By ChIP-seq, we observed a clear genome-wide correlation between the MLL-AF4 binding profile in *^CRISPR^MLL-AF4+* ALL, the *MLL-AF4* B-ALL SEM cell line^31^ and a primary *MLL-AF4* ALL patient sample^23^ (Supplementary Fig. 5d) and a substantial overlap in MLL-AF4 target genes (2,323 genes bound by MLL-AF4 in all 3 datasets) (Fig. 4c, Supplementary Table 4), with strikingly similar binding profiles, for example at *RUNX1* (Fig 4d).

**Fig. 4.**
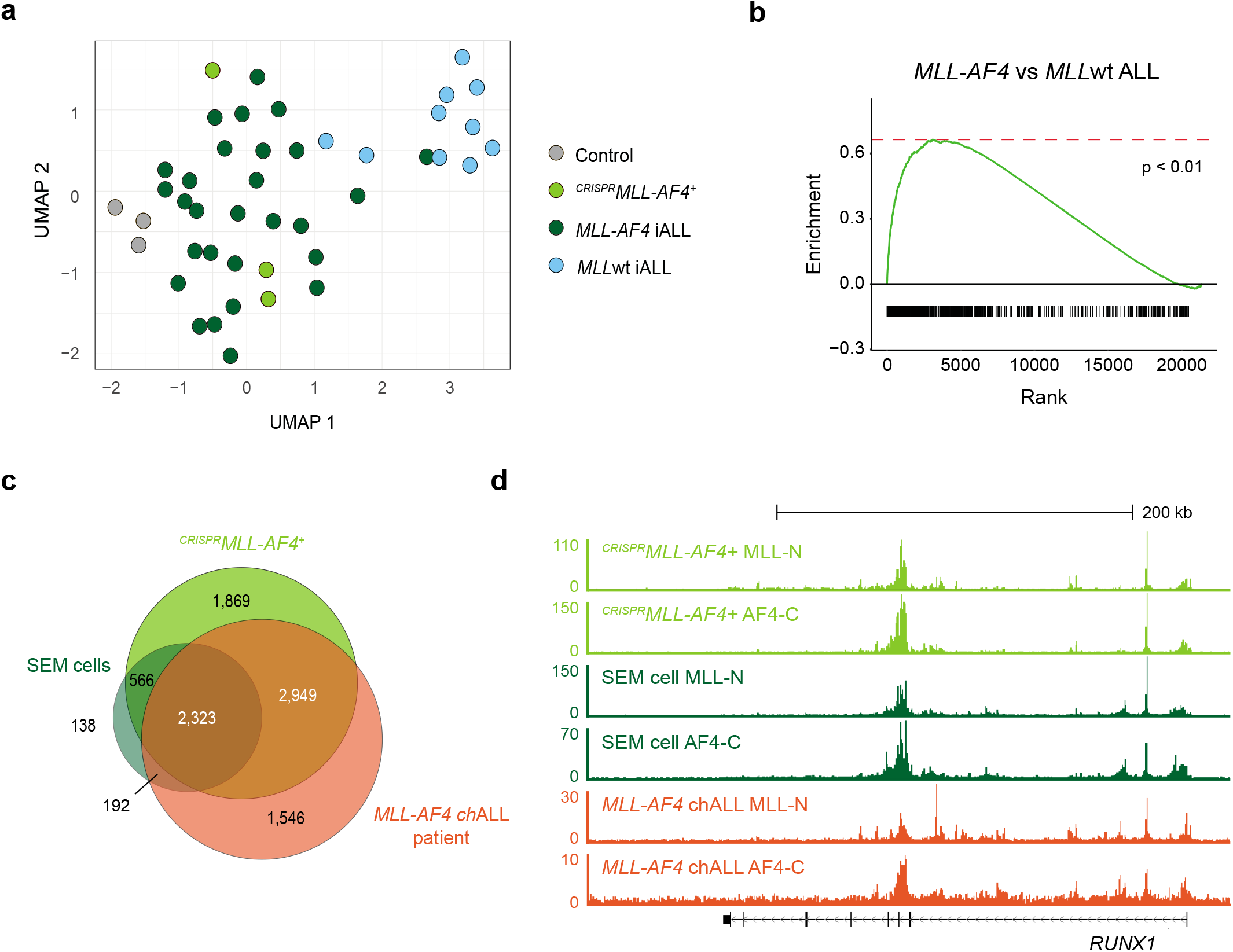
*^CRISPR^MLL-AF4+* ALL recapitulates the molecular profile of *MLL-AF4* ALL in patients. **a.** UMAP showing clustering of CD19+ cells from *^CRISPR^MLL-AF4^+^* and control mice with *MLL-AF4* (dark green) and *MLL*wt (blue) infant-ALL patient samples from a publicly available dataset ^19^ based on all differentially expressed genes (7,041 genes) between these 4 groups (edgeR generalized linear model (GLM), FDR < 0.05). **b.** Gene set enrichment analysis (GSEA) showing *^CRISPR^MLL-AF4+* ALL is more enriched for genes that are upregulated in *MLL-AF4* ALL compared to *MLLwt* ALL (1000 genes) when compared to CD19+ cells from control mice (p<0.01). **c.** Venn diagram showing overlap of MLL-AF4-bound genes (genes with an MLL-AF4 peak in the gene body) in *^CRISPR^MLL-AF4+* ALL in a primary recipient mouse, the *MLL-AF4+* SEM cell line and a primary *MLL-AF4* childhood-ALL (chALL) patient sample. MLL-AF4 peaks = directly overlapping MLL-N and AF4-C peaks. **d.** Representative ChIP-seq tracks at the MLL-AF4 target gene, *RUNX1* in *^CRISPR^MLL-AF4+* ALL, the SEM cell line and a primary *MLL-AF4* childhood-ALL (chALL) patient sample.

Finally, we wanted to ask whether inducing an *MLL-AF4* translocation in human FL gave rise to a model that specifically recapitulated the molecular profile of *MLL-AF4* infant-ALL. The only humanized mouse model of *MLL-AF4* ALL that has previously been published introduced a chimeric *MLL-Af4* fusion gene into human neonatal (cord blood (CB)) HSPC (hereafter referred to as CB *MLL-Af4+* ALL)^27^. We hypothesized that it may recapitulate *MLL-AF4* childhood-ALL, and could be used as a comparison to *^CRISPR^MLL-AF4+* ALL.

To examine the fetal and post-natal gene expression programs that are key to determining the age-related differences between *MLL-AF4* ALLs, we used the 139 genes up- or downregulated in both FL (compared to ABM) and *MLL-AF4* infant-ALL (compared to *MLL-AF4* childhood-ALL) (Supplementary Table 2). Clustering analysis based on this core gene list showed that, while *^CRISPR^MLL-AF4+* ALL was similar to *MLL-AF4* infant-ALL patients, CB *MLL-Af4+* ALL clustered away from *MLL-AF4* infant-ALL and closer to *MLL-AF4* childhood-ALL patients (Fig. 5a). To explore this in more detail, we carried out differential gene expression analysis between *^CRISPR^MLL-AF4+* ALL and CB *MLL-Af4+* ALL, followed by Gene Set Enrichment Analysis (GSEA). We found that *^CRISPR^MLL-AF4+* ALL was significantly enriched for genes upregulated in both FL HSPCs and *MLL-AF4* infant-ALL compared to CB *MLL-Af4+* ALL (Fig. 5b).

**Fig. 5.**
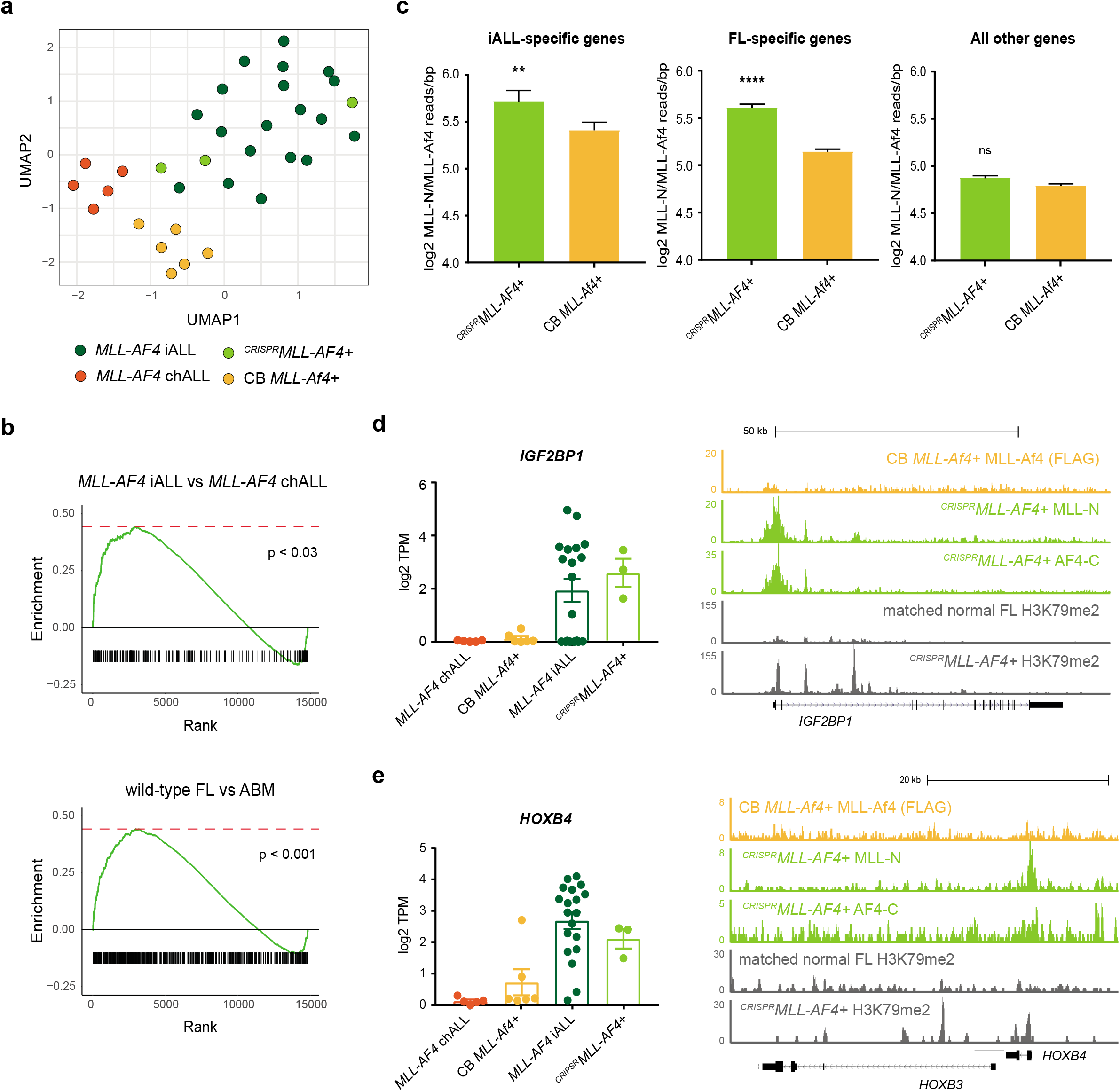
FL-derived *^CRISPR^MLL-AF4+* ALL specifically recapitulates the molecular profile of *MLL-AF4* infant-ALL (iALL) **a.** UMAP showing clustering of *^CRISPR^MLL-AF4+* ALL with *MLL-AF4* infant-ALL and away from *MLL-AF4* childhood-ALL (chALL) from a publicly available dataset^15^ and the CB *MLL-Af4+* ALL mouse model^27^ based on a set of genes differentially expressed in both FL (compared to ABM) and *MLL-AF4* infant-ALL (compared to *MLL-AF4* childhood-ALL) (139 genes, see Supplementary Table 2). **b.** Gene set enrichment analyses (GSEA) showing *^CRISPR^MLL-AF4+* ALL is significantly more enriched for genes that are upregulated in *MLL-AF4* infant-ALL (compared to *MLL-AF4* childhood-ALL) (617 genes, p<0.03) and in FL (compared to ABM) (5,709 genes, p<0.001) when compared to CB *MLL-Af4+* ALL. **c.** Bar plots showing MLL-AF4 enrichment (MLL-N ChIP-seq reads/bp normalized to 10^7^ total reads) is greater in *^CRISPR^MLL-AF4+* ALL than in CB *MLL-Af4+* ALL (FLAG ChIP-seq reads/bp normalized to 10^7^ total reads) at iALL-specific genes (193 genes, Mann-Whitney test, p<0.01) and FL-specific genes (3,949 genes, Mann-Whitney test, p<0.0001). Data shown as mean ± SEM. **d.** (left) Barplot showing expression of *IGF2BP1* in *MLL-AF4* childhood-ALL (chALL)^15^, CB *MLL-Af4+* ALL^27^, *MLL-AF4* infant-ALL (iALL)^15^ and *^CRISPR^MLL-AF4+* ALL. Data shown as mean ± SEM. (right) ChIP-seq tracks for *MLL-Af4* (FLAG ChIP-seq) in CB *MLL-Af4+* ALL (yellow), MLL-N and AF4-C ChIP-seq in *^CRISPR^MLL-AF4+* ALL and H3K79me2 ChIP-seq in *^CRISPR^MLL-AF4+* ALL and the biologically-matched unedited FL samples from which *^CRISPR^MLL-AF4+* ALL was derived, at *IGF2BP1*. ChIP-seq data shown is normalized to 10^7^ total reads. **e.** (left) Barplot showing expression of *HOXB4* in *MLL-AF4* childhood-ALL (chALL)^15^, CB *MLL-Af4+* ALL^27^, *MLL-AF4* infant-ALL (iALL)^15^ and *^CRISPR^MLL-AF4+* ALL. Data shown as mean ± SEM. (right) ChIP-seq tracks for *MLL-Af4* (FLAG ChIP-seq) in CB *MLL-Af4+* ALL (yellow), MLL-N and AF4-C ChIP-seq in *^CRISPR^MLL-AF4+* ALL and H3K79me2 ChIP-seq in *^CRISPR^MLL-AF4+* ALL and the biologically-matched unedited FL samples from which *^CRISPR^MLL-AF4+* ALL was derived, at *HOXB3/HOXB4*. ChIP-seq data shown is normalized to 10^7^ total reads.

Comparing MLL-AF4 binding at promoters genome-wide in both models, we found that MLL-AF4 in *^CRISPR^MLL-AF4+* ALL showed greater enrichment (normalized ChIP-seq reads/bp) at the promoters of infant-ALL- and FL-specific genes compared to MLL-Af4 in CB *MLL-Af4+* ALL (Fig. 5c). However, at all other genes, MLL-AF4/MLL-Af4 enrichment was comparable (Fig. 5c). At iALL- and FL-specific genes *IGF2BP1* (Fig. 5d) and *HOXB4* (Fig. 5e), we observed an MLL-AF4 peak in *^CRISPR^MLL-AF4+* ALL but not in CB *MLL-Af4+* ALL. These data suggest that MLL-AF4 may play an active role in maintaining fetal gene expression programs in infant-ALL. Increased levels of H3K79me2 are a commonly used marker of MLL-AF4 activity^23,28^. Therefore, using one of the unique features of our model, we carried out H3K79me2 ChIP-seq for the first time in identical primary human FL HSPC before and after leukemic transformation. We observed increased levels of H3K79me2 at MLL-AF4 peaks in FL- and iALL-specific genes such as *IGF2BP1* (Fig. 5d) and *HOXB4* (Fig. 5e) in *^CRISPR^MLL-AF4+* ALL, further suggesting that MLL-AF4 actively maintains the expression of these fetal-specific genes in *MLL-AF4* infant-ALL.

The mechanisms by which the same *MLL-r* driver mutation could cause more aggressive disease and worse outcomes in infant-ALL compared to childhood-ALL have always been unclear. We hypothesized that there must be intrinsic biological differences between infant-ALL and childhood-ALL blasts, unrelated to the driver mutation, that underlie these age-related differences. Here, we identify the unique molecular profile of *MLL-AF4* infant-ALL using primary patient data. Reasoning that this profile drives the distinct phenotype of infant-ALL, we set out to identify factors that could explain it. We find that maintenance of fetal-specific gene expression programs account for a large proportion (~40%) of the unique molecular profile of *MLL-AF4* infant-ALL, suggesting that it is the specific fetal target cell(s) in which it arises that provide the permissive cellular context for aggressive infant-ALL.

Human fetal HSPCs are more proliferative than ABM HSPCs^32,33^, and they differentiate down distinct developmental pathways^34,35^, some of which are virtually absent in adult life. Therefore, maintenance of fetal HSPC characteristics provides a possible explanation for the highly-proliferative, therapy-resistant nature of infant-ALL. One of the biggest challenges to understanding the biology of infant-ALL and developing novel, more effective therapies has been the lack of pre-clinical models^36^ that capture the unique characteristics and aggressive nature of the disease. By targeting a t(4;11)/*MLL-AF4* translocation to primary human FL HSPCs, we have created the first bona fide *MLL-AF4* infant-ALL model. Our results finally confirm that a human fetal cell context is permissive, and indeed probably required; to give rise to an ALL that recapitulates key phenotypic and molecular features of poor prognosis *MLL-AF4* infant-ALL.

*^CRISPR^MLL-AF4+* mice represent a previously lacking model in which the function of MLL-AF4 can be investigated in the appropriate human fetal cell context. Moreover, because *^CRISPR^MLL-AF4+* cells were generated by CRISPR-Cas9 genome editing, they express both *MLL-AF4* and the reciprocal *AF4-MLL* at physiological levels. Therefore, *^CRISPR^MLL-AF4+* ALL also provides an opportunity to explore the contribution of the reciprocal fusion protein during leukemogenesis, which has been a topic of debate in the *MLL-r* ALL field^37,38^. Finally, the infant-ALL-like features of *^CRISPR^MLL-AF4+* ALL make this an important model for future preclinical testing of novel therapies. To our knowledge, we are the first to report CNS disease in an *MLL-AF4* mouse model, which is a common clinical feature of infant-ALL that can lead to CNS relapse^4^. Therefore, the ability of novel treatments to eradicate blasts from the CNS is an important consideration, and this can now be tested in *^CRISPR^MLL-AF4+* ALL.

## ACKNOWLEDGEMENTS

T.A.M., S.R., N.T.C., and N.F. were funded by Medical Research Council (MRC, UK) Molecular Haematology Unit grant MC_UU_12009/6 and MC_UU_00016/6. S.O.B was funded by the Department of Paediatrics and Alexander Thatte Fund, University of Oxford. A.R. was supported by a Bloodwise Clinician Scientist Fellowship (grants: 14041 and 17001), Wellcome Trust Clinical Research Career Development Fellowship (216632/Z/19/Z), Lady Tata Memorial International Fellowship, and EHA-ASH Translational Research Training in Hematology Fellowship. I.R. is supported by the NIHR Oxford BRC, by a Bloodwise Program Grant (13001) and by the MRC Molecular Haematology Unit (MC_UU_12009/14). T.J. was supported as part of Wellcome Trust CRCDF (216632/Z/19/Z). C.H was funded by the Little Princess Trust & Children’s Cancer and Leukaemia group (CCLG 2017-13). N.E. was supported as part of Blood Cancer UK (1259; myeloid preleukaemia of Down syndrome). We gratefully acknowledge the Translational Histopathology Laboratory (THL) at the CRUK Oxford Centre, Department of Oncology for processing, sectioning and staining spleen and liver tissue samples and Lynn Stevenson and Clare Orange, University of Glasgow, for brain histology and imaging. We would also like to acknowledge the WIMM Flow Cytometry Facility which is supported by the MRC HIU; MRC MHU (MC_UU_12009); NIHR Oxford BRC; Kay Kendall Leukemia Fund (KKL1057), John Fell Fund (131/030 and 101/517), the EPA fund (CF182 and CF170) and by the WIMM Strategic Alliance awards G0902418 and MC_UU_12025. We thank the High-Throughput Genomics Group at the Wellcome Trust Centre for Human Genetics (funded by Wellcome Trust Grant Reference 090532/Z/ 09/Z); the MRC WIMM Centre for Computational Biology (CCB), Radcliffe Department of Medicine, University of Oxford; and Jelena Telenius for the use of her pipelines. The human fetal material was provided by the Joint MRC/Wellcome Trust Grant 099175/Z/ 12/Z Human Developmental Biology Resource (http://hdbr.org). We thank Prof Pablo Menendez for helpful advice, and gratefully acknowledge the kind generosity of patients, their parents and staff at Great Ormond Street Hospital, London.

## AUTHOR CONTRIBUTIONS

S.R., I.R., A.R., and T.A.M. conceived the experimental design; S.R., N.T.C., N.F., T.J., C.H., D.L., and S.O.B. carried out experiments; S.R., N.T.C, T.J., C.H., and A.R. analyzed and curated the data; S.R., C.H., I.R., A.R., and T.A.M. interpreted the data; S.R., I.R., A.R. and T.A.M. wrote the original manuscript; S.R., N.T.C., T.J., S.I., J.B., A.J.M., C.H., I.R., A.R., T.A.M. contributed to reviewing and editing the manuscript. I.R., A.R., and T.A.M. provided supervision and funding.

## ONLINE METHODS

### Fetal Samples

Donated fetal tissue was provided by the Human Developmental Biology Resource (HDBR, www.hdbr.org), regulated by the UK Human Tissue Authority (HTA, www.hta.gov.uk) and covered under ethics (REC: 18/NE/0290 and 18/LO/0822). FL samples used for CRISPR/Cas9 *MLL-AF4* translocation experiments underwent CD34 magnetic bead selection at the time of sample processing and were cryopreserved for future use as described previously^39^. MLL-AF4 ALL patient samples were obtained from Blood Cancer UK Childhood Leukaemia Cell Bank, UK (REC: 16/SW/0219). Patient samples were anonymized at source, assigned a unique study number and linked.

### Animals

All experiments were performed under a project license approved by the UK Home Office under the Animal (Scientific Procedures) Act 1986 and in accordance with the principles of 3Rs (replacement, reduction and refinement) in animal research.

### CRISPR-Cas9 *MLL-AF4* translocation

CRISPR-Cas9 genome editing was carried out using a previously described protocol^40^. *MLL* and *AF4* sgRNAs (Synthego) were first tested for editing efficiency individually in FL CD34+ cells. Cryopreserved CD34+ cells from a single primary human FL sample were thawed and placed into suspension culture at a density of 2.5×10^5^ cells/ml in StemLine II (Sigma) supplemented with SCF (100ng/ml), FLT3L (100ng/ml) and TPO (100ng/ml) (Peprotech) for 12 hours. Cells were harvested and electroporated with either (i) Cas9 protein (IDT) only or (ii) a Cas9/sgRNA RNP using a Neon™ Transfection System (Thermo Fisher). Electroporated cells were placed into fresh suspension culture media to recover overnight. Cells were harvested and bulk genomic DNA was extracted using a DNeasy Blood and Tissue Kit (Qiagen). A ~1 kb region of DNA around the target cut site was amplified by PCR and Sanger sequenced (Eurofins). Sanger sequencing traces from samples edited with RNPs were compared to traces from Cas9 only controls using the ICE Analysis online tool (Synthego, https://ice.synthego.com). Editing efficiency is reported as the percentage of indels detected (Supplementary Fig. 3a).

For each CRISPR-Cas9 *MLL-AF4* translocation experiment, cryopreserved CD34+ cells from a single 13-15 pcw primary human FL underwent suspension culture as described. Cells were harvested and electroporated with either (i) Cas9 protein (IDT) only, (ii) Cas9 protein plus *MLL*-sgRNA only, as biologically matched controls or (iii) a 1:1 mix of Cas9/MLL-sgRNA and Cas9/AF4-sgRNA RNPs using a Neon™ Transfection System (Thermo Fisher). Electroporated cells were placed into fresh suspension culture media to recover overnight before subsequent *in vitro* culture and *in vivo* transplantation experiments.

### MS-5 stroma co-culture

Electroporated FL CD34+ cells (*^CRISPR^MLL-AF4+* and control) were plated onto a confluent layer of MS-5 stromal cells in a 24-well plate at a density of 2,000 cells/well in αMEM (Gibco) supplemented with 10% heat-inactivated batch-tested FBS, 100U/ml Penicillin, 100μg/ml Streptomycin, 2mM L-glutamine, 50μM 2-Mercaptoethanol, 10mM HEPES, SCF (20ng/ml), FLT3L (10ng/ml), IL-2 (10ng/ml) and IL-7 (5ng/ml). Cultures were maintained as previously described^35,39^. Cells were harvested for flow cytometry analysis once a week beginning at week 2 of culture. *MLL-AF4* and *AF4-MLL* RT-qPCRs were carried out on week 4 of culture.

### Xenograft transplantation

8-12 week old female NSG mice were sub-lethally irradiated with two doses of 1.25Gy six hours apart (2.5Gy total) and injected via the tail vein with 25,000-35,000 edited FL cells (*^CRISPR^MLL-AF4+*, n=3; Cas9 control, n=5; or Cas9 plus *MLL*-sgRNA control, n=1) plus 30,000 wild-type, unedited, sex-mismatched FL CD34+ carrier cells. Engraftment was monitored by peripheral blood sampling every 3 weeks. Human CD45+ cells were sorted from peripheral blood samples to carry out *MLL-AF4* and *AF4-MLL* RT-qPCR for the detection of successfully edited cells. Animals were monitored regularly using a standardized physical scoring system, and any mouse found to be in distress was humanely killed. Mice were considered leukemic if they met at least 3 of the following criteria: (i) overt signs of disease (hunching, lack of movement, weight loss, paralysis), (ii) splenomegaly, (iii) PB blast count over 50%, (iv) peripheral organ infiltration, (v) detection of the *MLL-AF4* translocation in both BM and spleen.

### Flow cytometry

Cells were stained with fluorophore-conjugated monoclonal antibodies in PBS with 2% FBS and 1mM EDTA for 30 minutes and analyzed using BD LSR II or Fortessa X50 instruments. Antibodies used are detailed in Supplementary Table 5. Analysis was performed using FlowJo software where gates were set using unstained and fluorescence minus one controls.

### Histopathology

On termination, samples of ~0.5-1cm^2^ were taken from the spleen and liver of *^CRISPR^MLL-AF4^+^* and Cas9 control mice and fixed in 10% formaldehyde. After fixation, tissues were processed and paraffin embedded. 4μm paraffin sections we cut onto Superfrost Plus adhesive slides, VWR, Cat No 406/0179/00. Haematoxylin and Eosin (H&E) was performed using the Vector Laboratories H&E kit, Cat No 3502, as per their recommended protocol and mounted using Vectamount, Vector Laboratories, Cat No H5000-60.

Murine heads were decalcified and processed as previously described^41^. Following paraffin wax embedding, 2.5μm sections were cut onto Poly-L-silane coated slides and stained with Gill’s haematoxylin and Putt’s eosin (both made in house). Slides were imaged on a NanoZoomer Digital Pathology (NDP) slide scanner (Hamamatsu) and analyzed with NDP.view 2 software.

### RT-qPCR

Total RNA was extracted from cells using an RNeasy Micro Kit (Qiagen). cDNA was generated from polyA mRNA using a SuperScript III kit (Invitrogen). qPCR was carried out on cDNA using SYBRGreen master mix (Thermo Fisher) and a QuantStudio3 Real-Time PCR System (Thermo Fisher). For list of qPCR primers used see Supplementary Table 5.

### RNA-sequencing

Approximately 3×10^5^ CD45+CD19+ cells were sorted from the bone marrow of 3 primary *^CRISPR^MLL-AF4^+^* recipient mice and 3 control primary recipient mice (Cas9 control, n=2; Cas9 plus *MLL*-sgRNA, n=1). Total RNA was extracted using an RNeasy Mini Kit (Qiagen). Poly(A) purification was conducted using the NEB Poly(A) mRNA magnetic isolation module as per the manufacturer’s protocol. Library preparation was carried out using the Ultra II Directional RNA Library Prep Kit (NEB, E7765). RNA libraries were sequenced by paired-end sequencing using a 150 cycle high output kit on a Nextseq 500 (Illumina). RNA-seq protocols for sorted subpopulations of FL HSPC have been previously described in^19^.

### IgH rearrangement analysis

Samples were screened for IgH complete (VH-DH-JH) and IgH incomplete (DH-JH) rearrangements using BIOMED-2 protocols to detect clonality. DNA was extracted from cells from the bone marrow of 3 primary *^CRISPR^MLL-AF4^+^* recipient mice. IgH rearrangements were analyzed as described in^35^.

### ChIP-sequencing

The full protocol is described in^31^. In short, up to 5×10^7^ cells were sonicated (Covaris) following the manufacturer’s protocol and incubated with antibody overnight. Magnetic protein A and G beads (ThermoFisher Scientific) were used to isolate antibody-chromatin complexes. Antibodies used are detailed in Supplementary Table 5. Beads were washed three times using a solution of 50mM HEPES-KOH (pH7.6), 500mM LiCl, 1mM EDTA, 1% NP40 and 0.7% sodium deoxycholate and once with Tris-EDTA. Samples were eluted and Proteinase K/RNase A-treated. Samples were purified using a ChIP Clean and Concentrator kit (Zymo). DNA libraries were generated using the NEBnext Ultra DNA library preparation kit for Illumina (NEB). Libraries were sequenced by paired-end sequencing using a 75 cycle high output kit on a Nextseq 500 (Illumina).

### NGS analysis

For RNA-seq, following sequencing, QC analysis was conducted using the fastQC package (http://www.bioinformatics.babraham.ac.uk/projects/fastqc). Reads were mapped to the human genome assembly using STAR. The featureCounts function from the Subread package was used to quantify gene expression levels using standard parameters. This was used to identify differential gene expression globally using the edgeR package. Differential gene expression was defined by an adjusted p-value (FDR) of less than 0.05. Infant ALL RNA-seq datasets were analyzed as described previously^35^.

To derive a FL vs ABM gene signature, bulk RNA-seq for sorted subpopulations of FL HSPC^19^ were compared to matched sorted subpopulations of ABM HSPC^20^ (FL HSC vs adult BM HSC, FL MPP vs adult BM MPP, FL LMPP vs adult BM LMPP and FL committed B progenitors (CBP) vs adult BM CLP). Genes that were differentially expressed between FL and ABM in at least one matched HSPC subpopulation were included in the gene signature. Genes that showed a significant change in opposite directions in different HSPC subtypes (e.g. upregulated in FL HSC vs ABM HSC, but downregulated in FL LMPP vs ABM LMPP) or in the normal vs leukemic setting (e.g. upregulated in FL HSPC vs ABM HSPC, but downregulated in *MLL-AF4* infant-ALL vs *MLL-AF4* childhood-ALL) were filtered out of the gene signature to leave a total of 5,709 genes (Supplementary Table 2).

For ChIP-seq, quality control of FASTQ reads, alignment, PCR duplicate filtering, blacklisted region filtering and UCSC data hub generation was performed using an in-house pipeline (https://www.biorxiv.org/content/10.1101/393413v1) as described. The HOMER tool makeBigWig.pl command was used to generate bigwig files for visualization in UCSC, normalizing tag counts to tags per 1×10^7^. ChIP-seq peaks were called using the HOMER tool findPeaks.pl with ChIP input sample used to estimate background signal. Gene profiles were generated using the HOMER tool annotatePeaks.pl.

### Statistics

Two-tailed Mann-Whitney, Log-rank (Mantel-Cox) tests and ANOVA followed by multiple comparisons testing were used to compare experimental groups as indicated in the figure legends. Statistical analyses were performed using GraphPad Prism v7.00 or R v4.0.1. Data are expressed as mean ± SEM unless otherwise indicated.

### Data availability

Further information and requests for resources and reagents may be directed to and will be fulfilled by the corresponding authors, Dr Anindita Roy and Dr Thomas A Milne.

The accession number for the RNA-seq and ChIP-seq data generated during this study is NCBI GEO: XXXXX

## Code availability

ChIP-seq data were analyzed using an in-house pipeline (https://www.biorxiv.org/content/10.1101/393413v1).

---

**Supplementary Fig. 1.**
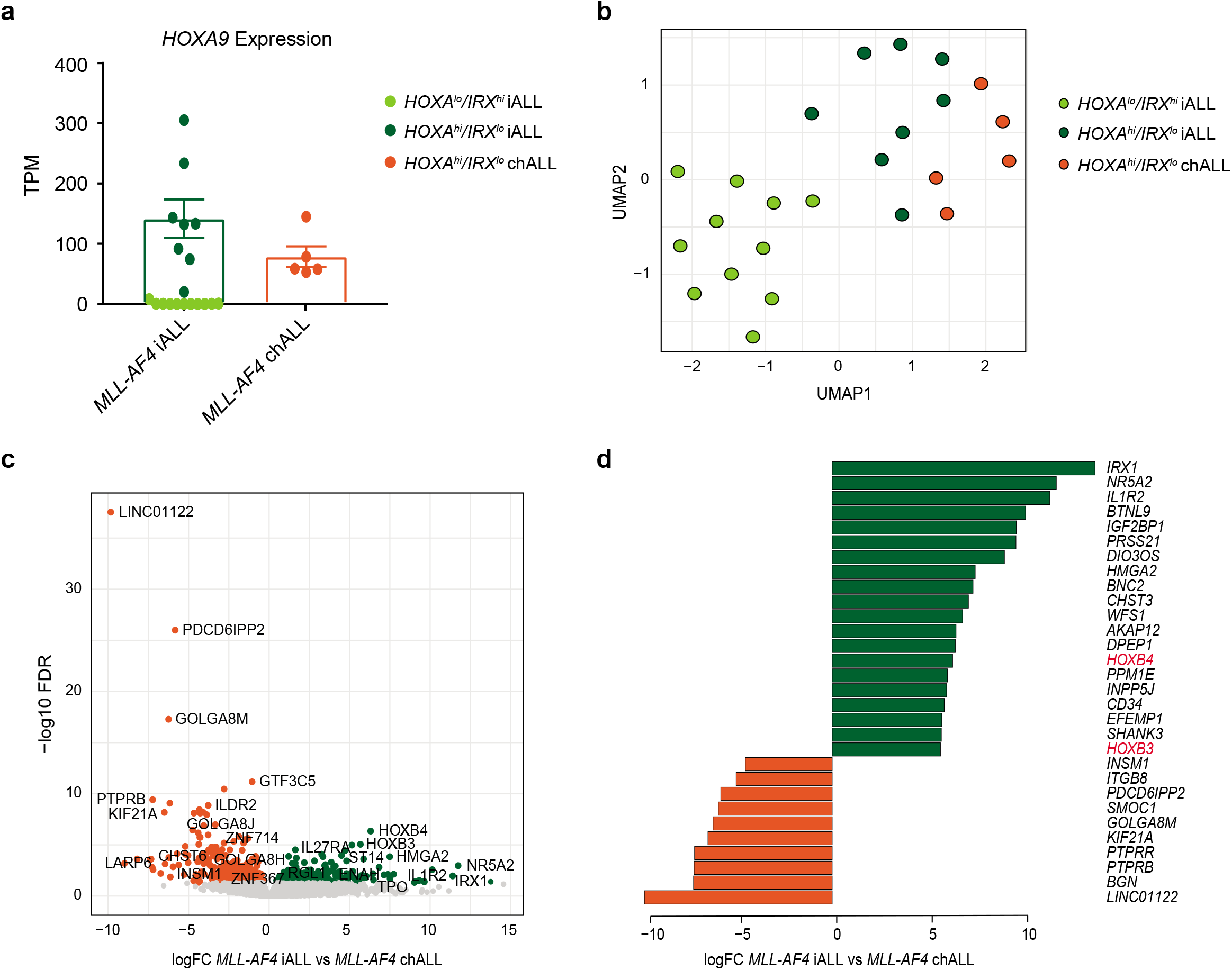
**a.** Barplot showing expression of *HOXA9* in *MLL-AF4* infant-ALL (iALL, n=19) and *MLL-AF4* childhood-ALL (chALL, n=5) from a previously published patient dataset^1^. Values are shown as transcripts per million (TPM). Data shown as mean ± SEM. Patients were considered to have a *HOXA^lo^/IRX^hi^* molecular profile when they showed a *HOXA9* expression < 20 TPM. (light green = *HOXA^lo^/IRX^hi^* infant-ALL (iALL), n=11; dark green = *HOXA^hi^/IRX^lo^* infant-ALL (iALL), n=8; orange = *HOXA^hi^/IRX^lo^* childhood-ALL (chALL)). No childhood-ALL patients in this dataset showed a *HOXA^lo^/IRX^hi^* molecular profile. **b.** UMAP showing clustering of *MLL-AF4* infant-ALL (*HOXA^lo^/IRX^hi^* iALL = light green, n=11; *HOXA^hi^/IRX^lo^* iALL = dark green, n=8) and *MLL-AF4* childhood-ALL (chALL = orange, n=5) from a previously published patient dataset^1^ based on the 500 most variable genes. **c.** Volcano plot showing all differentially expressed genes between MLL-AF4 infant-ALL and MLL-AF4 childhood ALL (dark green = significantly upregulated in MLL-AF4 infant-ALL (FDR,0.05, logFC>0); orange = significantly upregulated in MLL-AF4 childhood-ALL (FDR<0.05, logFC<0), gray = not significantly differentially expressed (FDR>0.05)). A selection of the most differentially expressed genes are labelled. **d.** Barplot showing the genes with the greatest logFC in *MLL-AF4* infant-ALL (green; top 20) and *MLL-AF4* childhood-ALL (orange; top 10).

**Supplementary Fig. 2.**
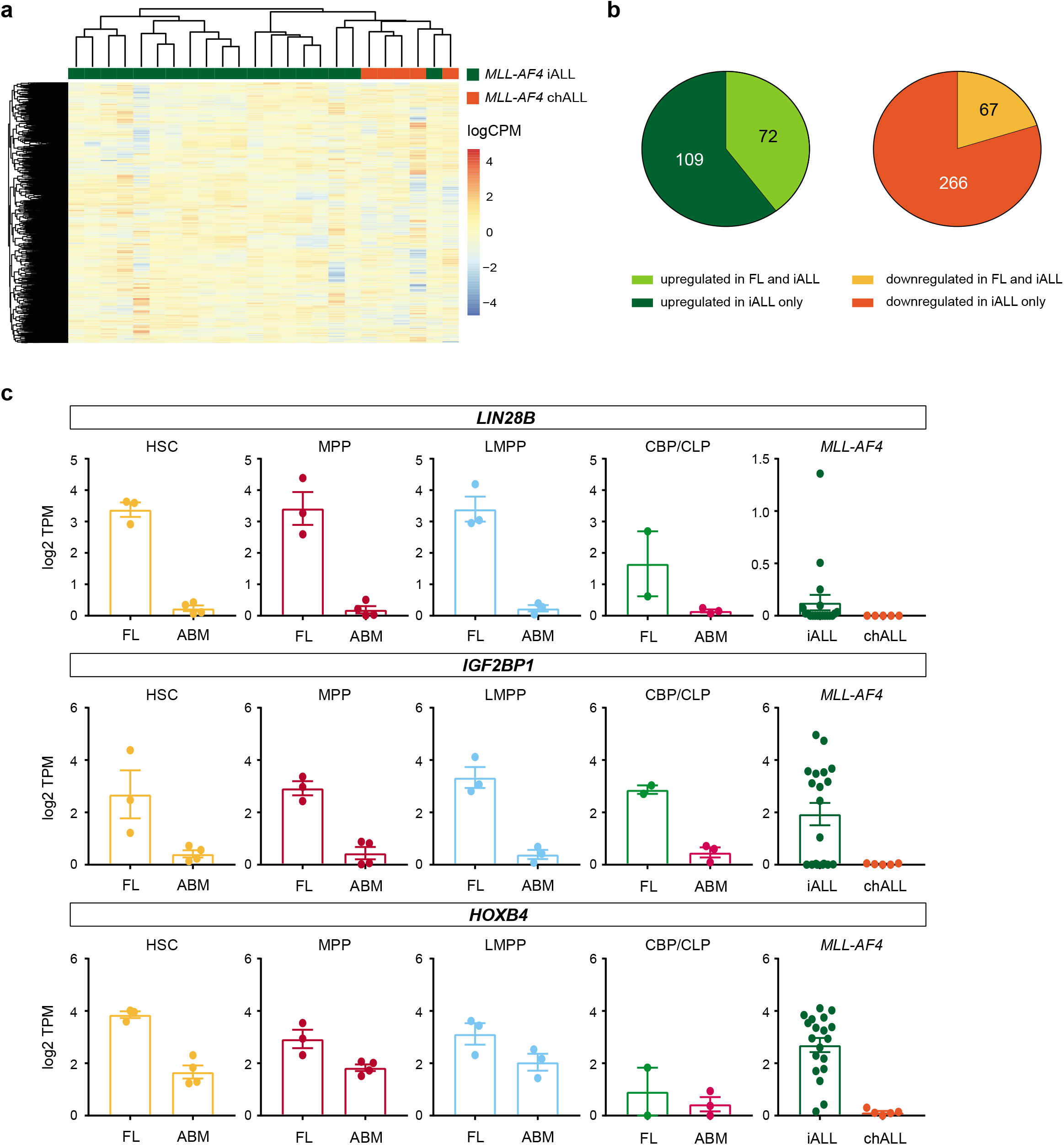
**a.** Heatmap showing clustering of *MLL-AF4* infant-ALL (iALL (green), n=19) and *MLL-AF4* childhood-ALL (chALL (orange), n=5) based on 5,709 significantly differentially expressed genes between FL and ABM HSPCs (FDR<0.05, Supplementary Table 2). Color scale = log2 counts per million (logCPM) **b.** Pie charts showing proportion of genes upregulated in *MLL-AF4* infant-ALL (compared to *MLL-AF4* childhood ALL; dark green) that are also upregulated in FL (compared to ABM; light green), and the proportion of genes downregulated in *MLL-AF4* infant-ALL (compared to *MLL-AF4* childhood ALL; orange) that are also downregulated in FL (compared to ABM; yellow) (see Supplementary Table 2). Values shown as number of genes. **c.** Barplots showing expression of *LIN28B, IGF2BP1* and *HOXB4* in FL and ABM HSPC subpopulations (HSC = hematopoietic stem cell, MPP = multipotent progenitor cell, LMPP = lymphoid-primed multipotent progenitor cell, CBP = committed B progenitor, CLP = common lymphoid progenitor), as well as *MLL-AF4* infant-ALL (iALL) and *MLL-AF4* chiildhood ALL (chALL). Data shown as mean ± SEM.

**Supplementary Fig. 3.**
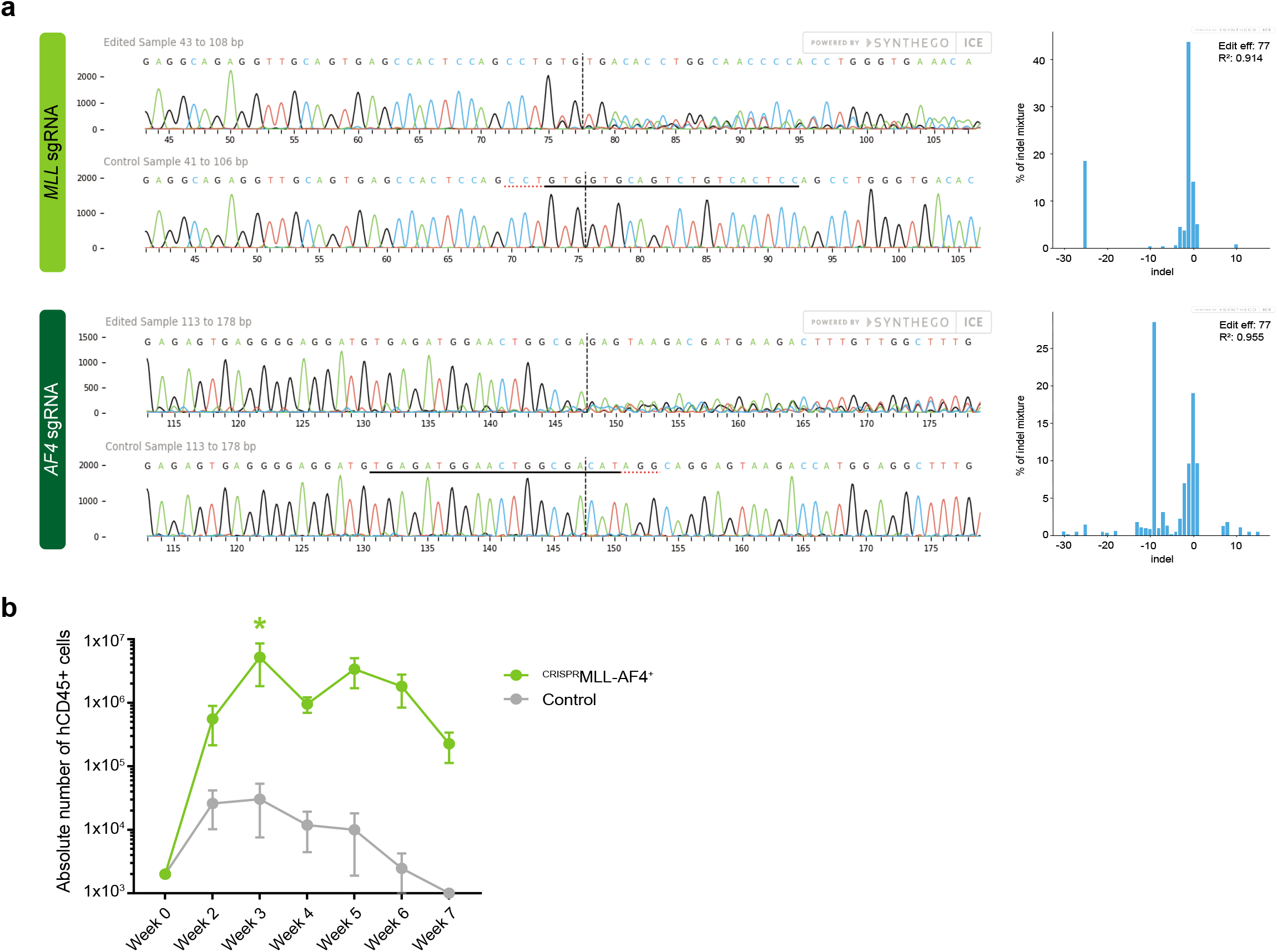
**a.** Synthego ICE Analysis (https://ice.synthego.com/) results for individual sgRNA efficiency tests for *MLL*-sgRNA and *AF4*-sgRNA in FL CD34+ cells. (left) Sanger sequencing tracks for edited cells (top) and unedited controls (bottom) around the PAM site. (right) Quantification of indels in edited cells. *MLL*-sgRNA and *AF4*-sgRNA both showed an editing efficiency of 77%. **b.** Cumulative absolute number of human CD45+ cells per well over time during MS-5 co-culture assay of *^CRISPR^MLL-AF4+* and control CD34+ cells (n=3). * =p<0.02 (Twoway ANOVA with Sidak correction for multiple comparisons). Data shown as mean ± SEM.

**Supplementary Fig. 4.**
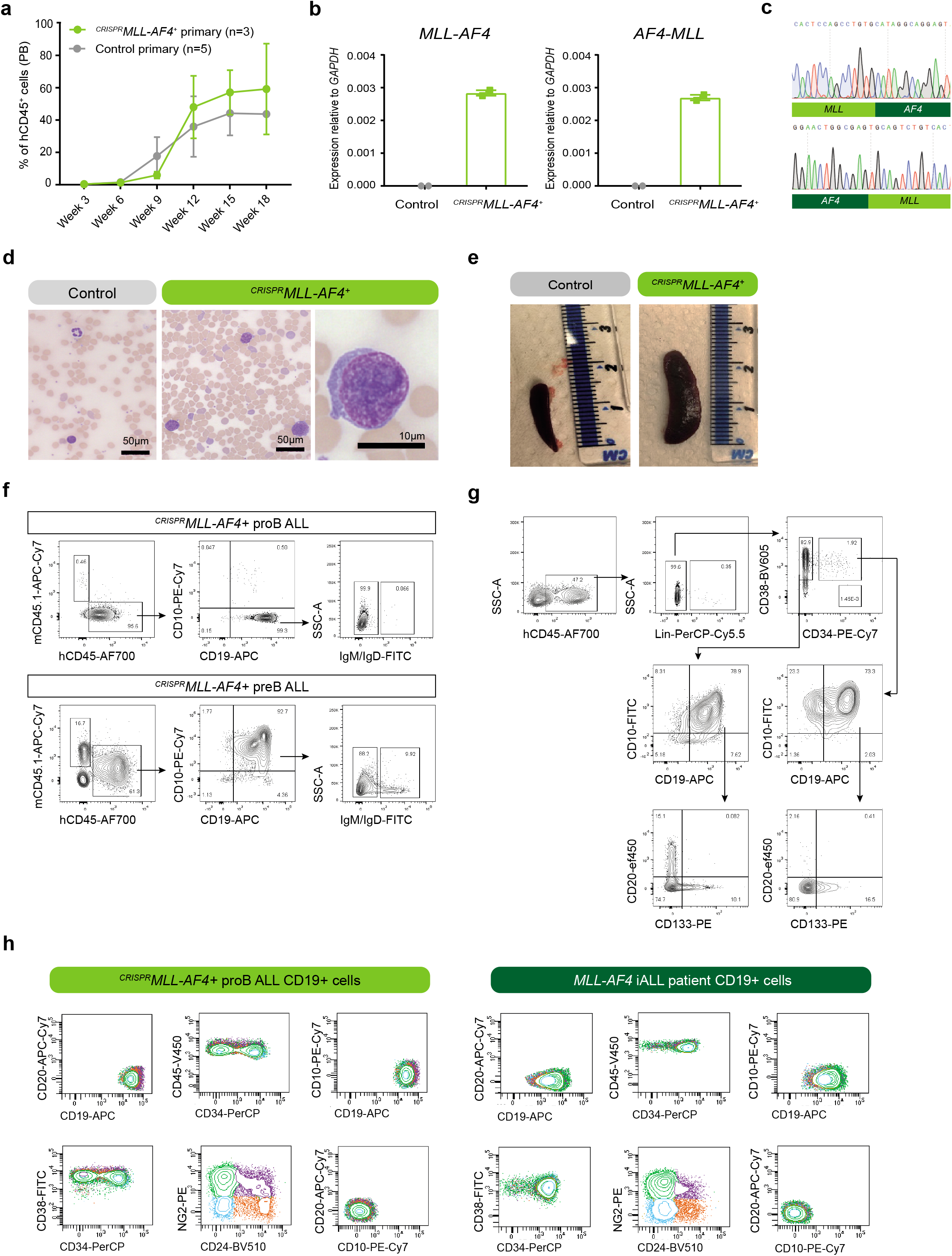
**a.** PB engraftment of human CD45+ (hCD45+) cells over time in primary *^CRISPR^MLL-AF4+* (n=3) and control (n=5) recipient mice. Quantified as a percentage of all CD45+ cells (mouse CD45.1+ and human CD45+). Data shown as mean ± SEM. **b.** RT-qPCR showing expression of *MLL-AF4* (n=2) and *AF4-MLL* (n=2) relative to *GAPDH* in human CD45+ cells isolated from PB at week 12 post-engraftment. **c.** Sanger sequencing tracks showing *MLL-AF4* and *AF4-MLL* genomic DNA breakpoints in blasts isolated from the spleen of *^CRISPR^MLL-AF4+* mice. Breakpoint regions were amplified by PCR before Sanger sequencing in order to examine the translocated allele without contamination by the remaining WT allele. *MLL* and *AF4* portions are labelled below each track. **d.** Representative H&E-stained peripheral blood (PB) films for control (left) and *^CRISPR^MLL-AF4+* (right) primary recipient mice. Low magnification images (scale bar = 50μm) show multi-lineage cells in the PB in controls and predominantly circulating blast cells in *^CRISPR^MLL-AF4+* mice. High magnification image (scale bar = 10μm) shows a representative blast cell from *^CRISPR^MLL-AF4+* PB. **e.** Representative images of the spleens of control and *^CRISPR^MLL-AF4+* mice. **f.** Representative flow cytometry plots of viable, single cells in proB *^CRISPR^MLL-AF4+* (top) and preB *^CRISPR^MLL-AF4+* (bottom) BM at termination. (mCD45.1, mouse CD45; hCD45, human CD45). **g.** Representative flow cytometry plots of viable, single cells in control and *^CRISPR^MLL-AF4+* BM at termination (week 17). **h.** Representative flow cytometry plots of CD19+ blasts from *^CRISPR^MLL-AF4+* BM at termination (week 18) (left) and an *MLL-AF4* iALL patient BM (right). Datapoints are colored in all plots based on surface NG2 and CD24 expression.

**Supplementary Fig. 5.**
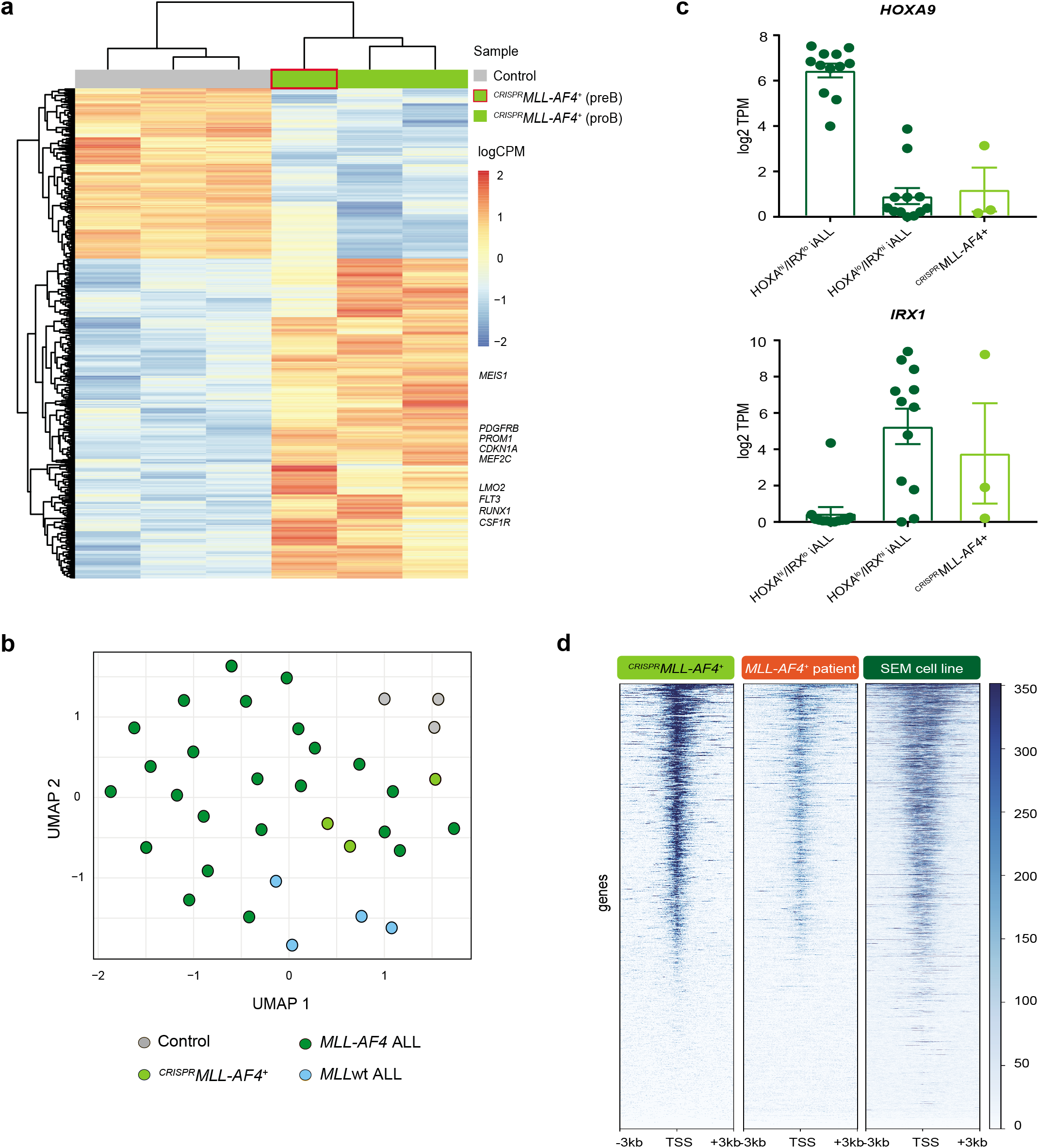
a. Heatmap showing significantly differentially expressed (FDR < 0.05) genes between primary control and *^CRISPR^MLL-AF4+* mice. A selection of genes known to be upregulated in *MLL-AF4* ALL are labelled. b. UMAP showing clustering of *^CRISPR^MLL-AF4+* (light green) and control (gray) mice with *MLL-AF4* (dark green) and *MLL*wt (blue) ALL patients from a publicly available dataset^1^ based on all differentially expressed genes (5,785 genes) between these 4 groups (edgeR generalized linear model (GLM)). c. Barplot showing expression of *HOXA9* and *IRX1* for *HOXA^hi^/IRX^lo^* infant-ALL (iALL), *HOXA^lo^/IRX^hi^* infant-ALL (iALL) and *^CRISPR^MLL-AF4+* ALL (designated *HOXA^lo^/IRX^hi^* based on *HOXA9* TPM <20). Data shown as mean ± SEM. d. Heatmap showing MLL-N ChIP-seq enrichment for a 6kb region centered on the promoter (transcriptional start site (TSS)) of all genes in *^CRISPR^MLL-AF4+* ALL, the SEM cell line and a primary *MLL-AF4* childhood-ALL (chALL) patient sample, sorted by MLL-N ChIP-seq signal in *^CRISPR^MLL-AF4+* ALL. Scale = reads/bp/10^7^ total reads.

## REFERENCES

1 Vora, A. et al. Treatment reduction for children and young adults with low-risk acute lymphoblastic leukaemia defined by minimal residual disease (UKALL 2003): a randomised controlled trial. Lancet Oncol 14, 199–209, doi:10.1016/S1470-2045(12)70600-9 (2013).

2 Pieters, R. et al. Outcome of Infants Younger Than 1 Year With Acute Lymphoblastic Leukemia Treated With the Interfant-06 Protocol: Results From an International Phase III Randomized Study. J Clin Oncol 37, 2246–2256, doi:10.1200/JCO.19.00261 (2019).

3 Moorman, A. V. et al. Prognostic effect of chromosomal abnormalities in childhood B-cell precursor acute lymphoblastic leukaemia: results from the UK Medical Research Council ALL97/99 randomised trial. The Lancet Oncology 11, 429–438, doi:10.1016/s1470-2045(10)70066-8 (2010).

4 Pui, C.-H. et al. Outcome of treatment in childhood acute lymphoblastic leukaemia with rearrangements of the 11q23 chromosomal region. The Lancet 359, 1909–1915, doi:10.1016/s0140-6736(02)08782-2 (2002).

5 Biondi, A., Cimino, G., Pieters, R. & Pui, C. H. Biological and therapeutic aspects of infant leukemia. Blood 96, 24–33 (2000).

6 Hilden, J. M. et al. Analysis of prognostic factors of acute lymphoblastic leukemia in infants: report on CCG 1953 from the Children’s Oncology Group. Blood 108, 441–451, doi:10.1182/blood-2005-07-3011 (2006).

7 Pieters, R. et al. A treatment protocol for infants younger than 1 year with acute lymphoblastic leukaemia (Interfant-99): an observational study and a multicentre randomised trial. Lancet 370, 240–250, doi:10.1016/S0140-6736(07)61126-X (2007).

8 Meyer, C. et al. The MLL recombinome of acute leukemias in 2017. Leukemia 32, 273–284, doi:10.1038/leu.2017.213 (2018).

9 Takahashi, S. & Yokoyama, A. The molecular functions of common and atypical MLL fusion protein complexes. Biochim Biophys Acta Gene Regul Mech 1863, 194548, doi:10.1016/j.bbagrm.2020.194548 (2020).

10 Iacobucci, I. & Mullighan, C. G. Genetic Basis of Acute Lymphoblastic Leukemia. J Clin Oncol 35, 975–983, doi:10.1200/JCO.2016.70.7836 (2017).

11 Mann, G. et al. Improved outcome with hematopoietic stem cell transplantation in a poor prognostic subgroup of infants with mixed-lineage-leukemia (MLL)-rearranged acute lymphoblastic leukemia: results from the Interfant-99 Study. Blood 116, 2644–2650, doi:10.1182/blood-2010-03-273532 (2010).

12 Tomizawa, D. et al. A risk-stratified therapy for infants with acute lymphoblastic leukemia: a report from the JPLSG MLL-10 trial. Blood 136, 1813–1823, doi:10.1182/blood.2019004741 (2020).

13 Reichel, M. et al. Biased distribution of chromosomal breakpoints involving the MLL gene in infants versus children and adults with t(4;11) ALL. Oncogene 20, 2900–2907, doi:10.1038/sj.onc.1204401 (2001).

14 Trentin, L. et al. Two independent gene signatures in pediatric t(4;11) acute lymphoblastic leukemia patients. Eur J Haematol 83, 406–419, doi:10.1111/j.1600-0609.2009.01305.x (2009).

15 Andersson, A. K. et al. The landscape of somatic mutations in infant MLL-rearranged acute lymphoblastic leukemias. Nat Genet 47, 330–337, doi:10.1038/ng.3230 (2015).

16 Greaves, M. In utero origins of childhood leukaemia. Early Hum Dev 81, 123–129, doi:10.1016/j.earlhumdev.2004.10.004 (2005).

17 Mullighan, C. G. et al. Genome-wide analysis of genetic alterations in acute lymphoblastic leukaemia. Nature 446, 758–764, doi:10.1038/nature05690 (2007).

18 Wiemels, J. L. et al. Prenatal origin of acute lymphoblastic leukaemia in children. Lancet 354, 1499–1503, doi:10.1016/s0140-6736(99)09403-9 (1999).

19 Agraz-Doblas, A. et al. Unravelling the cellular origin and clinical prognostic markers of infant B-cell acute lymphoblastic leukemia using genome-wide analysis. Haematologica, doi:10.3324/haematol.2018.206375 (2019).

20 Corces, M. R. et al. Lineage-specific and single-cell chromatin accessibility charts human hematopoiesis and leukemia evolution. Nat Genet 48, 1193–1203, doi:10.1038/ng.3646 (2016).

21 Copley, M. R. et al. The Lin28b-let-7-Hmga2 axis determines the higher self-renewal potential of fetal haematopoietic stem cells. Nat Cell Biol 15, 916–925, doi:10.1038/ncb2783 (2013).

22 Elcheva, I. A. et al. RNA-binding protein IGF2BP1 maintains leukemia stem cell properties by regulating HOXB4, MYB, and ALDH1A1. Leukemia 34, 1354–1363, doi:10.1038/s41375-019-0656-9 (2020).

23 Godfrey, L. et al. H3K79me2/3 controls enhancer-promoter interactions and activation of the pan-cancer stem cell marker PROM1/CD133 in MLL-AF4 leukemia cells. Leukemia, doi:10.1038/s41375-020-0808-y (2020).

24 Pui, C. H. Acute lymphoblastic leukemia in children. Curr Opin Oncol 12, 3–12, doi:10.1097/00001622-200001000-00002 (2000).

25 Roberts, I., Fordham, N. J., Rao, A. & Bain, B. J. Neonatal leukaemia. Br J Haematol 182, 170–184, doi:10.1111/bjh.15246 (2018).

26 Silverman, L. B. Acute lymphoblastic leukemia in infancy. Pediatr Blood Cancer 49, 1070–1073, doi:10.1002/pbc.21352 (2007).

27 Lin, S. et al. Instructive Role of MLL-Fusion Proteins Revealed by a Model of t(4;11) Pro-B Acute Lymphoblastic Leukemia. Cancer Cell 30, 737–749, doi:10.1016/j.ccell.2016.10.008 (2016).

28 Krivtsov, A. V. et al. H3K79 methylation profiles define murine and human MLL-AF4 leukemias. Cancer Cell 14, 355–368, doi:10.1016/j.ccr.2008.10.001 (2008).

29 Metzler, M. et al. A conditional model of MLL-AF4 B-cell tumourigenesis using invertor technology. Oncogene 25, 3093–3103, doi:10.1038/sj.onc.1209636 (2006).

30 Lopez-Millan, B. et al. NG2 antigen is a therapeutic target for MLL-rearranged B-cell acute lymphoblastic leukemia. Leukemia, doi:10.1038/s41375-018-0353-0 (2019).

31 Kerry, J. et al. MLL-AF4 Spreading Identifies Binding Sites that Are Distinct from Super-Enhancers and that Govern Sensitivity to DOT1L Inhibition in Leukemia. Cell Rep 18, 482–495, doi:10.1016/j.celrep.2016.12.054 (2017).

32 Lansdorp, P. M., Dragowska, W. & Mayani, H. Ontogeny-related changes in proliferative potential of human hematopoietic cells. J Exp Med 178, 787–791, doi:10.1084/jem.178.3.787 (1993).

33 Rebel, V. I., Miller, C. L., Eaves, C. J. & Lansdorp, P. M. The repopulation potential of fetal liver hematopoietic stem cells in mice exceeds that of their liver adult bone marrow counterparts. Blood 87, 3500–3507 (1996).

34 Notta, F. et al. Distinct routes of lineage development reshape the human blood hierarchy across ontogeny. Science 351, aab2116, doi:10.1126/science.aab2116 (2016).

35 O’Byrne, S. et al. Discovery of a CD10 negative B-progenitor in human fetal life identifies unique ontogeny-related developmental programs. Blood, doi:10.1182/blood.2019001289 (2019).

36 Milne, T. A. Mouse models of MLL leukemia: recapitulating the human disease. Blood 129, 2217–2223, doi:10.1182/blood-2016-10-691428 (2017).

37 Kumar, A. R., Yao, Q., Li, Q., Sam, T. A. & Kersey, J. H. t(4;11) leukemias display addiction to MLL-AF4 but not to AF4-MLL. Leuk Res 35, 305–309, doi:10.1016/j.leukres.2010.08.011 (2011).

38 Bursen, A. et al. The AF4.MLL fusion protein is capable of inducing ALL in mice without requirement of MLL.AF4. Blood 115, 3570–3579, doi:10.1182/blood-2009-06-229542 (2010).

39 Roy, A. et al. Perturbation of fetal liver hematopoietic stem and progenitor cell development by trisomy 21. Proc Natl Acad Sci U S A 109, 17579–17584, doi:10.1073/pnas.1211405109 (2012).

40 Gundry, M. C. et al. Highly Efficient Genome Editing of Murine and Human Hematopoietic Progenitor Cells by CRISPR/Cas9. Cell Rep 17, 1453–1461, doi:10.1016/j.celrep.2016.09.092 (2016).

41 Shi, Y. et al. Phase II-like murine trial identifies synergy between dexamethasone and dasatinib in T-cell acute lymphoblastic leukemia. Haematologica, doi:10.3324/haematol.2019.241026 (2020).

## REFERENCES

1 Andersson, A. K. et al. The landscape of somatic mutations in infant MLL-rearranged acute lymphoblastic leukemias. Nat Genet 47, 330–337, doi:10.1038/ng.3230 (2015).

